# The small molecule regacin binds a conserved pocket in RegA to inhibit the expression of bacterial virulence factors

**DOI:** 10.64898/2026.09.11.750636

**Authors:** Chris M Bollinger, Grace V Lawhern, Kacey M Talbot, F. Jon Kull, Charles R Midgett, George P Munson

**Affiliations:** Department of Microbiology & Immunology, University of Miami, Leonard M. Miller School of Medicine, Miami, FL 33101; Department of Chemistry, Dartmouth College, Hanover, NH 03755

## Abstract

Widespread antibiotic resistance is an urgent threat to public health. Small molecules that specifically target virulence factors or their expression are promising alternative therapeutics to antibiotics. The small molecule regacin was identified as a potent inhibitor of RegA, a transcription factor required for the expression of numerous virulence genes of *Citrobacter rodentium.* This demonstrates that targeting virulence regulators is a viable strategy for developing clinically useful therapeutics. RegA is a member of the AraC/XylS superfamily of transcription factors and with homologs in numerous pathogens that contribute to human morbidity and mortality worldwide. The development of antivirulence therapeutics that target these regulators could be accelerated by understanding the structural basis of inhibition. To that end we present the first high resolution structural analysis of RegA with and without bound regacin. RegA is structurally homologous to ToxT of *Vibrio cholerae* and Rns of enterotoxigenic *Escherichia coli.* Regacin was bound within a conserved binding pocket within the RegA amino-terminal domain. Structure guided mutagenesis of key pocket residues confirmed their role in regacin mediated inhibition. These findings define the regacin binding site, the structural basis for RegA inhibition, and provide a framework for designing inhibitors that target a vast family of virulence regulators essential for the pathogenesis of numerous bacterial pathogens.

## Introduction

Antimicrobial resistance is a public health crisis driven by poor drug stewardship and exacerbated by the private sector’s retreat –and in many cases, abandonment– of antibiotic research and development due to economic disincentives (Plackett, 2020). Antivirulence therapeutics are a promising alternative to traditional antibiotics because they specifically target virulence factors or inhibit their expression. Due to this specificity, there is less pressure for the development of resistance to these therapeutics (Dickey et al., 2017). In addition, they preserve the host microbiome because they are hyper-narrow spectrum often targeting a single species or even a single pathotype within a species. The preservation of the microbiome protects the host from opportunistic pathogens that take advantage of antibiotic driven dysbiosis (Rafey et al., 2023). This is another advantage of this class of therapeutics.

A handful of small molecule antivirulence therapeutics have advanced into clinical trials. ALS-4 has completed Phase I trials. It inhibits the synthesis of staphyloxanthin rendering *Staphylococcus aureus* –including MRSA– susceptible to reactive oxygen species generated by immune cells of the host (Walesch et al., 2022). Clinical trials are also underway for two small molecules that target the fimbrial adhesins of uropathgogenic *Escherichia coli* and enteropathogenic spp. of Enterobacteriaceae (Heimann et al., 2025; Reinisch et al., 2020) Promising preclinical antivirulence small molecules include LED209 that targets QseC, a histidine sensor kinase, to inhibit the expression of virulence factors in several Gram-negative species (Curtis et al., 2014; Rasko et al., 2008). M21 renders *S. aureus* avirulent by targeting the ClpP protease to downregulate alpha-hemolysin and a quorum-sensing system (Gao et al., 2018). Savirin disrupts the expression of many of the same virulence factors as M21 but does not target the ClpP protease. Rather, Savirin specifically binds to and inhibits the transcription factor AgrA drives the expression of alpha hemolysin and other virulence factors (Sully et al., 2014).

Antivirulence proof-of-concept has also been demonstrated by targeting RegA, the central virulence regulator of the murine enteropathogen *Citrobacter rodentium*. The expression of numerous *C. rodentium* virulence factors is dependent on RegA, and *regA* mutants are attenuated in murine infection models (Hart et al., 2008). This murine pathogen is often used to model human enteric infections because it shares many virulence factors in common with host restricted human diarrheal pathogens including enterohemorrhagic, enteropathogenic, enteroaggregative, and enterotoxigenic *E. coli* (ETEC). The discovery and development of antivirulence drugs against these enteropathogens is important because diarrheal disease is the second leading cause of death in children under five years of age in low resource nations (Fekadu et al., 2025).

RegA is a member of the AraC/XylS superfamily of transcription factors defined by a DNA-binding domain (DBD) containing two helix-turn-helix motifs (Gallegos et al., 1997; Yang et al., 2011). The DBD of most superfamily members is often accompanied by an amino-terminal domain responsible for dimerization and ligand-binding (Lowden et al., 2010; Midgett et al., 2021; Schüller et al., 2012; Soisson et al., 1997). To identify RegA inhibitors Yang et al. developed a Lac reporter strain to screen a library of ca. 12,000 small molecules (Yang et al., 2013). The best lead compound (CAS 345993-15-9) of the first screen was then used to seed a computational search of a vastly larger library of nearly 1 million molecules. Their secondary search led to the identification of a more potent analog, regacin (5-[(4-chloro-benzyl)thio]-1,3,4-thiadiazol-2-amine; CAS 72836-33-0). As expected of an antivirulence molecule, regacin did not inhibit the growth of *C. rodentium* or *E. coli*. Cytotoxicity was also found to be negligible against an immortalized mammalian cell line. In vivo, orally administered regacin significantly reduced the ability of *C. rodentium* to colonize the intestines of mice (Yang et al., 2013). To better understand the mechanism(s) of antivirulence drugs that inhibit AraC/XylS family members we present the first high resolution structures of apo and regacin bound RegA and structure guided genetic analysis of the regacin binding pocket.

## Results

### Identification of a ligand binding pocket within RegA

The apo-RegA crystal structure was solved to 2.69 Å revealing four structurally similar monomers making up two homodimers (average pairwise RMSD of 0.91 ± 0.17 Å) (**Fig. 1A and 1C**). The most notable difference among the four chains is the presence of an additional 10 residues at the amino-terminal end of chain A, which is disordered in the other chains (**Fig. 1B**). The overall structure and domain organization of apo-RegA is similar to our previously reported structures of *Vibrio cholerae* ToxT and Rns from ETEC (Lowden et al., 2010; Midgett et al., 2021). Like those AraC/XylS virulence regulators, the carboxy-terminal DBD of RegA is composed of seven α-helices with dual helix-turn-helix motifs that are the signature of all AraC/XylS family members. The RegA amino-terminal domain is composed of dimerization helices and a ligand binding pocket within a cupin fold (**Fig. 1C**). While the overall shape of the binding pocket differs in ToxT, Rns, and RegA, all three have polar residues lining the entrance and hydrophobic residues lining the interior (**Fig. 2**).

**Figure 1:**
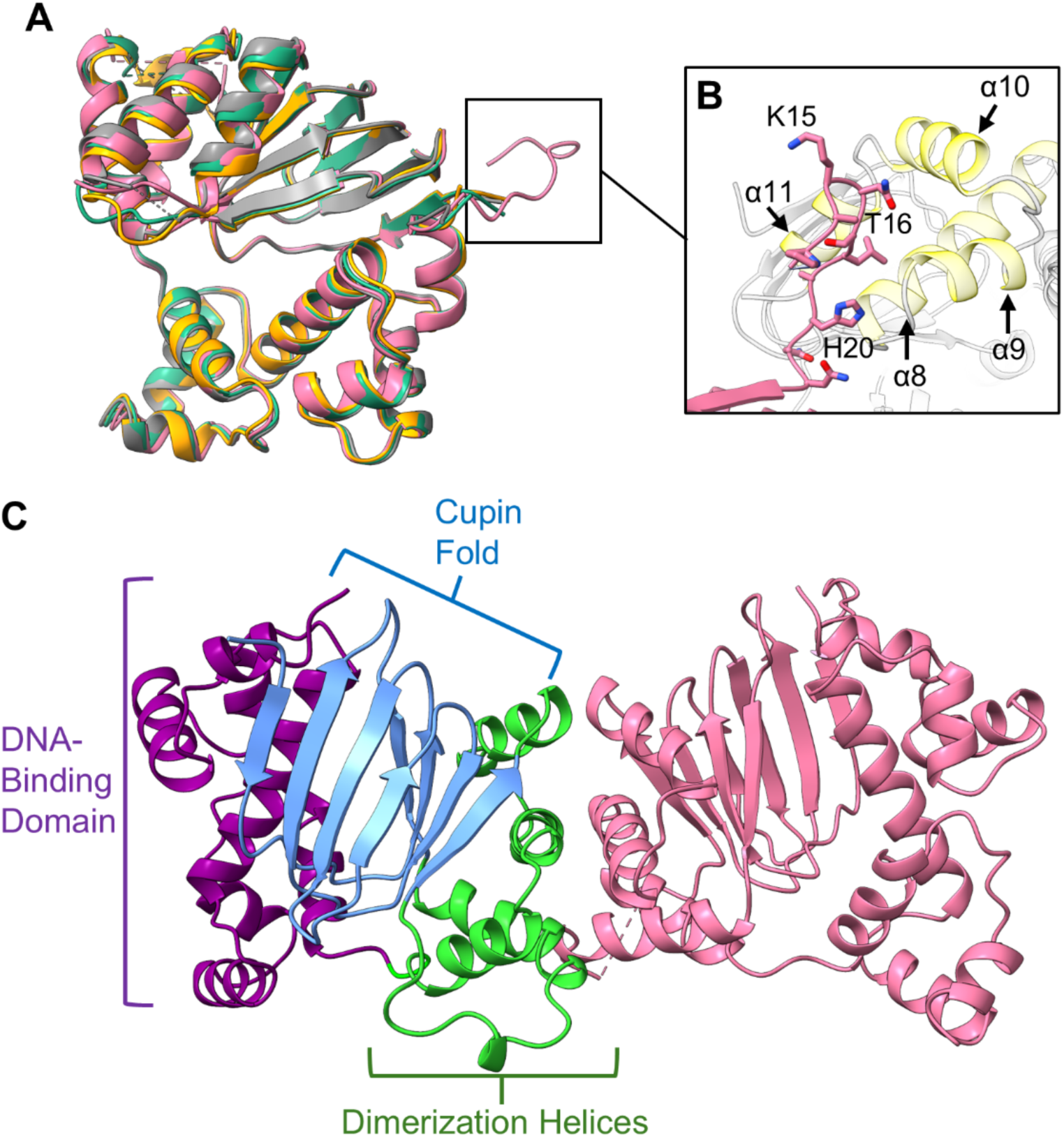
RegA is crystallized as a dimer. **A)** RegA crystallizes as a dimer at 2.69 Å resolution (PDB_00001873). Each monomer in the apo-RegA asymmetric unit is aligned, and they are nearly superimposable. Chain A is shown in pink, B is orange, C is teal and D is gray. Chain breaks are shown as dashed lines. **B)** RegA chain A amino-terminus (pink) interacting with the symmetry mate DBD helices α8, α9, α10, and α11 (yellow). Amino acids H20, T16, and K15 are indicated as reference markers. **C)** RegA Chain A monomer is shown in pink, while Chain B of the dimer is colored to indicate structure features. The dimerization interface helices (green) and cupin fold (cornflower blue) make up the amino-terminal domain. The DNA binding domain (purple) is indicated, and loops between are shown in gray.

**Figure 2:**
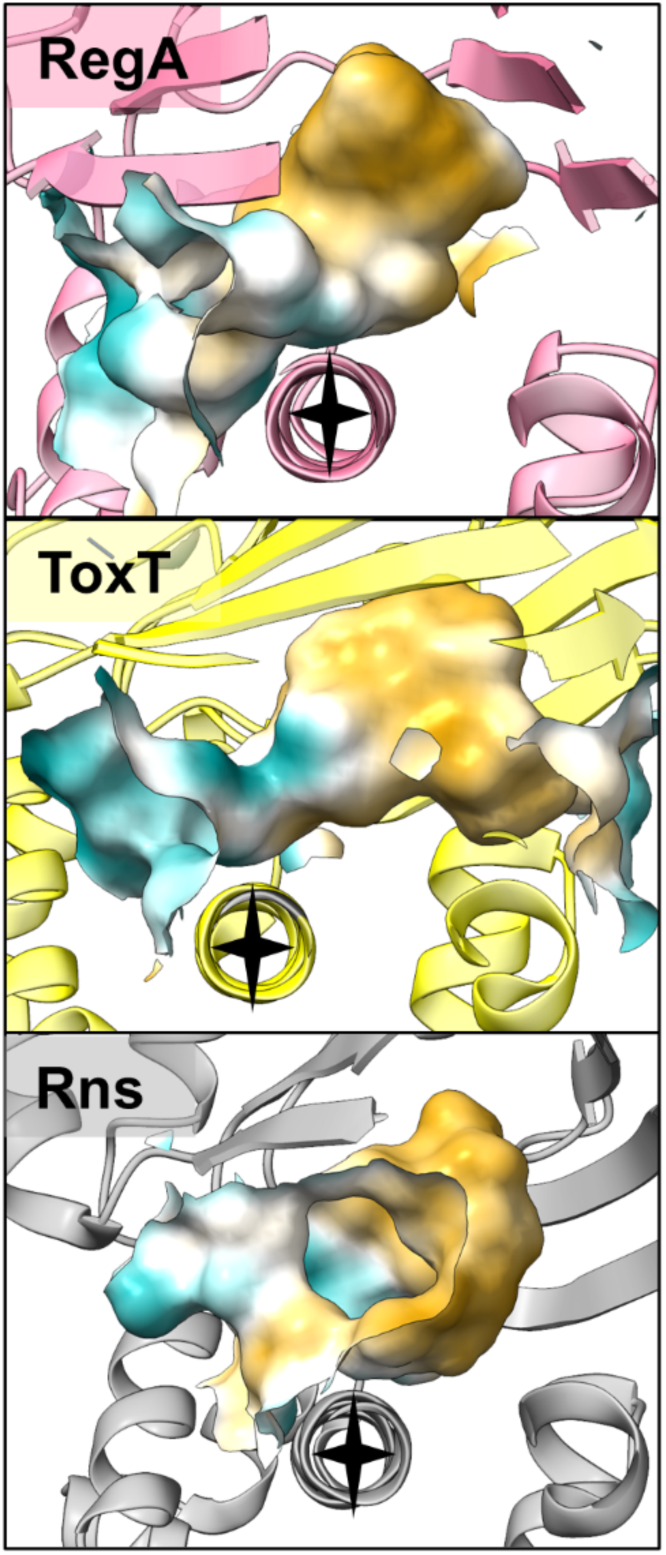
RegA homologs share ligand binding pocket surface topologies. Homologs ToxT (PDB ID: 3GBG, yellow) and Rns (PDB ID: 6XIU, gray) are aligned to RegA chain A (PDB_00001873, pink) α8 long helix (residues 215-233). The long helix for all structures is indicated by a black star. Pocket surfaces are defined as a 7 Å zone around the ligand bound within each pocket, and the surface is colored by hydrophobicity. Blue denotes polar surfaces and orange denotes hydrophobic surfaces.

### Regacin binds within the RegA ligand binding pocket

Regacin abolishes RegA DNA binding, and it has been proposed that the most likely binding site is within the DBD (Yang et al., 2013). To determine how RegA interacts with regacin, we solved the structure of RegA in the presence of regacin. Four monomers are arranged to form two dimers within the asymmetric unit as seen in apo-RegA with both binding pockets of one of the dimers occupied by regacin. Like the apo-RegA structure, regacin-RegA crystallized in the same space group with comparable unit cell dimensions (**Table S1**). The amino-terminal end of chain A was ordered, making similar contacts with a symmetry mate as in the apo-RegA structure (**Fig. S1**). Therefore, the unit cell consists of a regacin bound dimer and an apo-dimer. Notably, ligand binding does not appear to alter the structure of RegA (**Fig. S2**). The apo-RegA dimer and the regacin-bound dimer are nearly superimposable (**Fig. 3A**). Furthermore, the positions of the interacting residue side chains of the ligand binding pocket do not show any major changes when compared to chain A of the apo-RegA structure (**Fig. 3B and 3C**).

**Figure 3:**
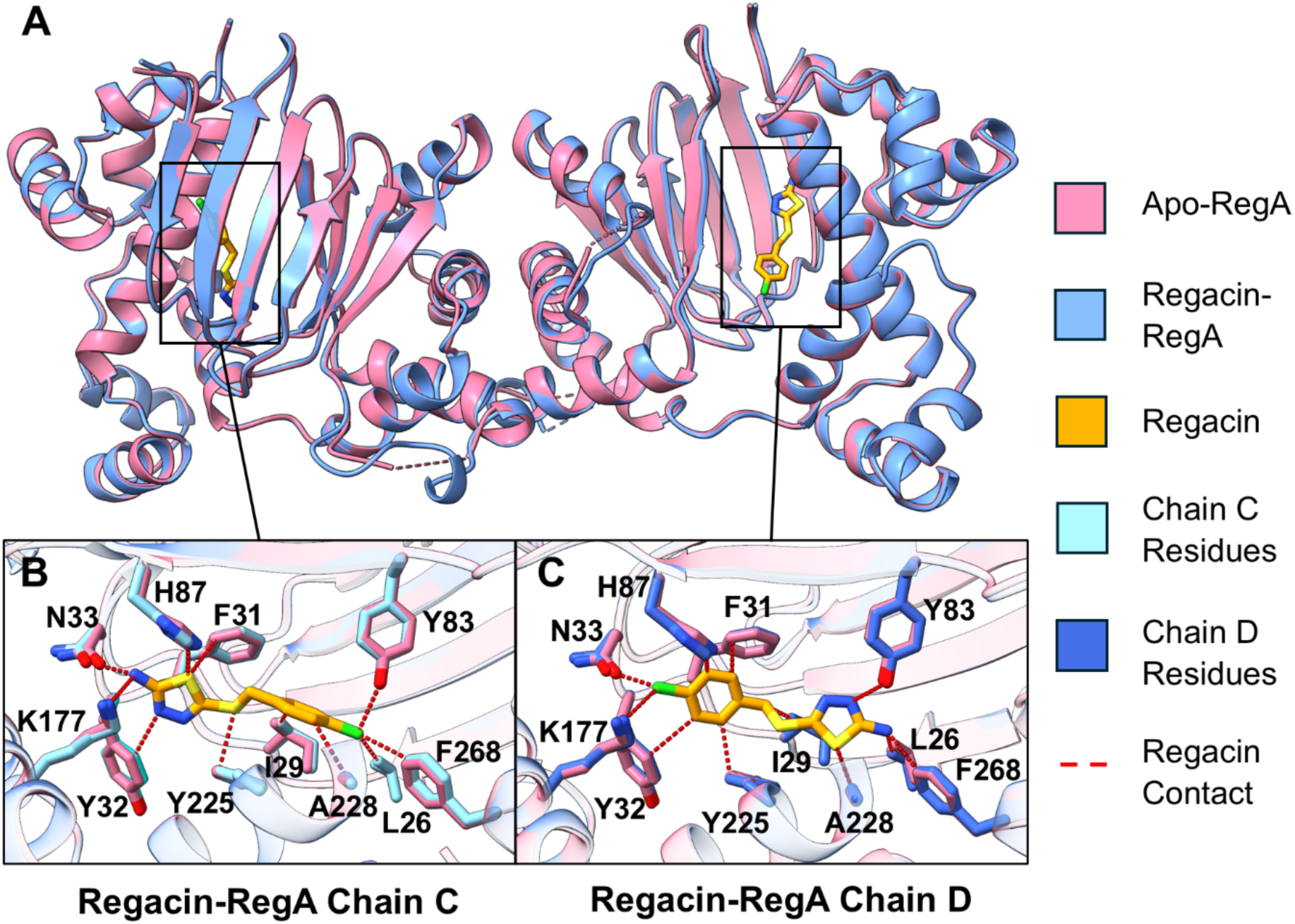
Apo-and regacin-bound RegA structures show no major structural changes or changes in binding pocket side chain orientation. **A)** Overlaid regacin-RegA chains C and D (PDB_00001874, blue) with apo-RegA chains C and D (PDB_00001873, pink). Regacin (orange) side chain contacts with distances ≤ 4 Å are shown in red. **B)** Side chain-regacin contacts are shown for regacin-RegA chain C (light blue) and the corresponding apo-RegA chain C (pink) side chains are shown. Contacts ≤ 4 Å between regacin (orange) and regacin-RegA are shown in red for chain C. **C)** Side chain-regacin contacts are shown for regacin-RegA chain D (dark blue) and the corresponding apo-RegA chain D (pink) side chains are shown Contacts ≤ 4 Å between regacin (orange) and regacin-RegA are shown in red for chain D.

Due to the pseudosymmetry of the regacin molecule and the resolution of the structure, orienting regacin within the pocket was challenging. As a result, composite omit maps and electron density fit validation metrics were used to place regacin in the two monomers with two different orientations related by a 180° rotation (**Fig. S3)**. In chain C, residues Y32, N33, and H87 are involved in hydrogen bonding with the 1,3,4-thiadiazole group of regacin. Several additional residues, including L26, I29, F31, Y83, K177, Y225, A228, and F268 contribute to hydrophobic and van der Waals contacts surrounding the molecule’s periphery (**Fig. 3B and Table 1**). In chain D, regacin forms a halogen bond between its chlorine atom and residue N33, and hydrophobic interactions with F31 and Y225. Residues L26, I29, Y32, Y83, H87, K177, A228, and F268 establish numerous van der Waals contacts at the periphery of the molecule (**Fig. 3C and Table 1**). Overall, side chains within the pocket interact with regacin in different but overlapping ways for each ligand orientation.

**Table 1:** Regacin-RegA residue level interactions, interaction types assessed with PLIP.

| Chain C |  |  | Chain D |  |
| --- | --- | --- | --- | --- |
| Residue | Interaction Type | Distance (Å) | Interaction Type | Distance (Å) |
| L26 | Van der Waals contact | 4.07 | Van der Waals contact | 3.19 |
| I29 | Hydrophobic Interaction | 3.94 | Van der Waals contact | 3.70 |
| F31 | Van der Waals contact | 4.07 | Hydrophobic Interaction | 3.38 |
| Y32 | Hydrogen Bond | 2.93 | Van der Waals contact | 4.18 |
| N33 | Hydrogen Bond | 2.79 | Halogen Bond | 3.00 |
| Y83 | Van der Waals contact | 3.22 | Van der Waals contact | 3.35 |
| H87 | Hydrogen Bond | 3.16 | Van der Waals contact | 3.08 |
| K177 | Van der Waals contact | 3.36 | Van der Waals contact | 3.25 |
| Y225 | Van der Waals contact | 3.52 | Hydrophobic Interaction | 3.29 |
| A228 | Hydrophobic Interaction | 3.78 | Van der Waals contact | 3.57 |
| F268 | Van der Waals contact | 3.35 | Van der Waals contact | 3.14 |

Hydrophobic InteractionHydrogen BondHalogen BondVan der Waals contact

### The RegA binding pocket exhibits plasticity

To elucidate which RegA residues contribute to regacin inhibition, we mutagenized and evaluated them for their ability to activate the CS3 pilin promoter (**Fig. 4A**) (Midgett et al., 2021). Some substitutions resulted in no transcriptional activation and were therefore excluded from further analysis (F31A, F31E, Y83A/V85A, V85F, Y86A, F261A). Polar residues that were mutagenized singly (Y32A, N33A, H87A, K177A, Y225A, and Y225E) displayed a range of regacin sensitivity. N33, H87, and K177 are likely to participate in electrostatic or hydrogen bonding contacts at the front of the binding pocket, while residue Y225 interacts from its position off the long helix of the DBD (**Fig. 4B**). Two double substitutions of interacting residues (H87A/K177A and H87A/Y225A) also failed to completely abolish regacin sensitivity. However, K177A/Y225A and H87A/K177A/Y225A both abolished ligand-induced inhibition (**Fig. 4A**). These data demonstrate that the RegA binding pocket makes non-specific polar contacts with regacin.

**Figure 4:**
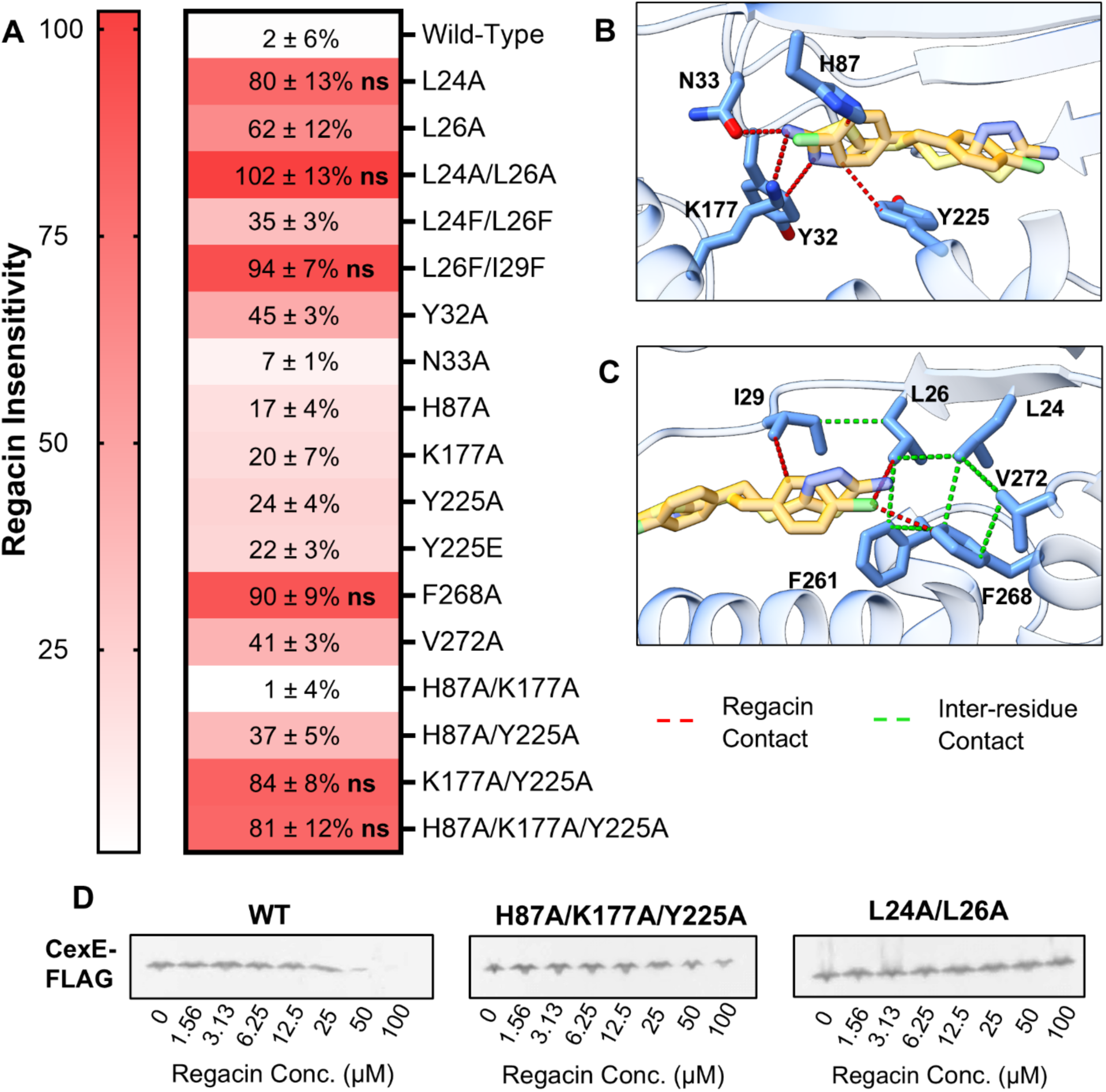
Pocket mutations show binding pocket plasticity. **A)** Heat map indicates disrupted residues and the impact on regacin sensitivity. “Regacin Insensitivity” refers to percent activity in CS3*p* reporter GPM1072 when grown in 24 μM regacin as compared to DMSO control (0.4% v/v DMSO). Data is given as percent activity compared to DMSO control ± SE. Residue modifications which show no statistical difference between DMSO and regacin treatment are denoted with “ns.” All residues not labeled “ns” were significantly inhibited by 24 μM regacin treatment. n = 3, ns *p-value* > 0.05 **B)** Contacts ≤ 4 Å between regacin (red orange) and polar front of pocket residues (blue) are shown in red while secondary hydrophobic contacts are shown in lime for regacin-RegA chain C (PDB_00001874). **C)** Contacts ≤4 Å between regacin (red orange) and hydrophobic back of pocket residues (blue) are shown in red while secondary hydrophobic contacts are shown in lime. **D)** Representative Western blots of *C. rodentium* DBS100 *cexE-FLAG/ΔregA/*pGPMRegA-myc, H87A/K177A/Y225A, and L24A/L26A cultured in the presence of regacin (1% v/v DMSO) probed with an anti-FLAG antibody.

At the back of the RegA binding pocket, there is a network of hydrophobic residues which bridge the cupin fold of the amino-terminal domain and the DBD (including L24, L26, I29, F261, F269, and V272), some of which regacin contacts (L26, I29, and F268) **(Fig. 4C).** L24A and F268A variants were insensitive to regacin, L26A resulted in greatly reduced regacin sensitivity, and V272A was only slightly less sensitive to regacin (**Fig. 4A**). Previous work in Rns identified two residues, I14 and I17, that sterically disrupt fatty acid binding when both were substituted to phenylalanine. To emulate this, the congruent RegA residues L26 and I29 were altered (Tolbert et al., 2025). When both L26 and I29 were simultaneously replaced with phenylalanine, RegA was insensitive to regacin. Altering both L24 and L26 did not have the same effect; RegA L24F/L26F was only roughly 30% less sensitive to regacin. In contrast to a space-filling construct, when L24 and L26 were both replaced with alanine, RegA became fully insensitive to regacin (**Fig. 4A**). Residue L26 contacts regacin within the co-crystal structure while L24 does not, indicating that perhaps this observed insensitivity is not a result of disrupted binding, but a result of disrupting broader hydrophobic contacts.

Assays measuring RegA variant activity are performed in an *E. coli* Lac reporter, so the function of two regacin-insensitive variants was observed qualitatively *in situ* in *C. rodentium* strain GPM3202 (*ΔregA*, *cexE-FLAG*). GPM3202 expressing the RegA-dependent virulence factor CexE-FLAG was complemented with RegA-WT, H87A/K177A/Y225A, and L24A/L26A constitutively expressive from a plasmid. Strain GPM3202 transformed with wild-type RegA, the triple elimination variant, or the hydrophobic elimination variant L24A/L26A were grown in the presence of various concentrations of regacin and whole cell lysates were stained for CexE-FLAG expression in Western blots (**Fig. 4D**). The H87A/K177A/Y225A variant is visibly less sensitive to regacin as compared to wild-type RegA, and CexE-FLAG expression was wholly unaffected by regacin when under the control of RegA-L24A/L26A. In summary, multiple eliminations of regacin contact residues in the front of the ligand binding pocket need to be substituted to fully abolish ligand-sensitivity. Beyond the front of the ligand binding pocket, several key residues in the hydrophobic back of the ligand binding pocket are also critical for ligand sensitivity.

## Discussion

The exquisite specificity of antivirulence drugs may limit or retard the development of resistance to them. Thus, the identification and development of such therapeutics has been proposed as a potential solution to the scourge of antimicrobial resistance that has rendered many antibiotics clinically ineffective (Sack et al., 1997; Salam et al., 2023; Wallace et al., 2020). Developed as a proof of concept, regacin (CAS 72836-33-0) has been shown to have several properties of an ideal antivirulence drug (Yang et al., 2013). First, it inhibits or abolishes the expression of RegA-dependent genes with negligible off target effects. Second, regacin does not inhibit the growth of *C. rodentium* or *E. coli*. Third, cytotoxicity was not observed when HeLa cells were cultured with regacin. Finally, in a murine colonization experiment orally administered regacin caused wild-type *C*. *rodentium* to phenocopy a *regA* mutant (Yang et al., 2013).

Yang et al. also investigated the molecular mechanism of regacin (Yang et al., 2013). Consistent with our homodimeric regacin-RegA structure (**Fig. 1 and 3**), they found that regacin does not disrupt RegA dimers by analytical ultracentrifugation. Regacin does, however, abolish the transcription factor’s ability to bind DNA as determined by EMSAs. When we compared our apo and regacin bound structures we found that the structures were essentially identical, lacking obvious differences that could promote or exclude DNA binding (**Fig. 3**). This is consistent with our prior studies of ToxT and Rns (Lowden et al., 2010; Midgett et al., 2021). In those prior studies we concluded that ligand binding does not produce large allosteric changes. Rather, ligand binding shifts the structural dynamics of each regulator. In the absence of ligand the regulators are flexible and this dynamism is essential for DNA binding. Ligand binding drives the regulators to a more rigid state incapable of binding DNA. Our analyses of apo and regacin bound structures suggest that this model of dynamic allostery is applicable to RegA.

Our structural analysis of the regacin binding pocket revealed an interior lined with hydrophobic residues and an entrance lined with polar residues. Although there are differences between them; the ligand binding pockets of RegA, Rns and ToxT have similar hydrophobic topologies (**Fig. 2**). Despite this similarity, ToxT is refractory to inhibition by regacin. In contrast, the antivirulence molecule significantly inhibits the activity of Rns (Yang et al., 2013). Our structural and genetic analyses of RegA indicate that its binding pocket can accommodate regacin in at least two orientations. Moreover, ligand binding is dependent upon a variety of hydrophobic and electrostatic interactions as well as hydrogen bonding. Previously, Yang et al. deployed random mutagenesis in an attempt to define the regacin binding site within RegA. They identified a single mutation, I222T, that renders RegA less sensitive to regacin. This mutation lies within the long helix that connects the dual helix-turn-helix motifs of the DBD. Although I222T is near the binding pocket, it projects away from it (**Fig. 5**). It is not understood how the I222T substitution abolishes regacin sensitivity.

**Figure 5:**
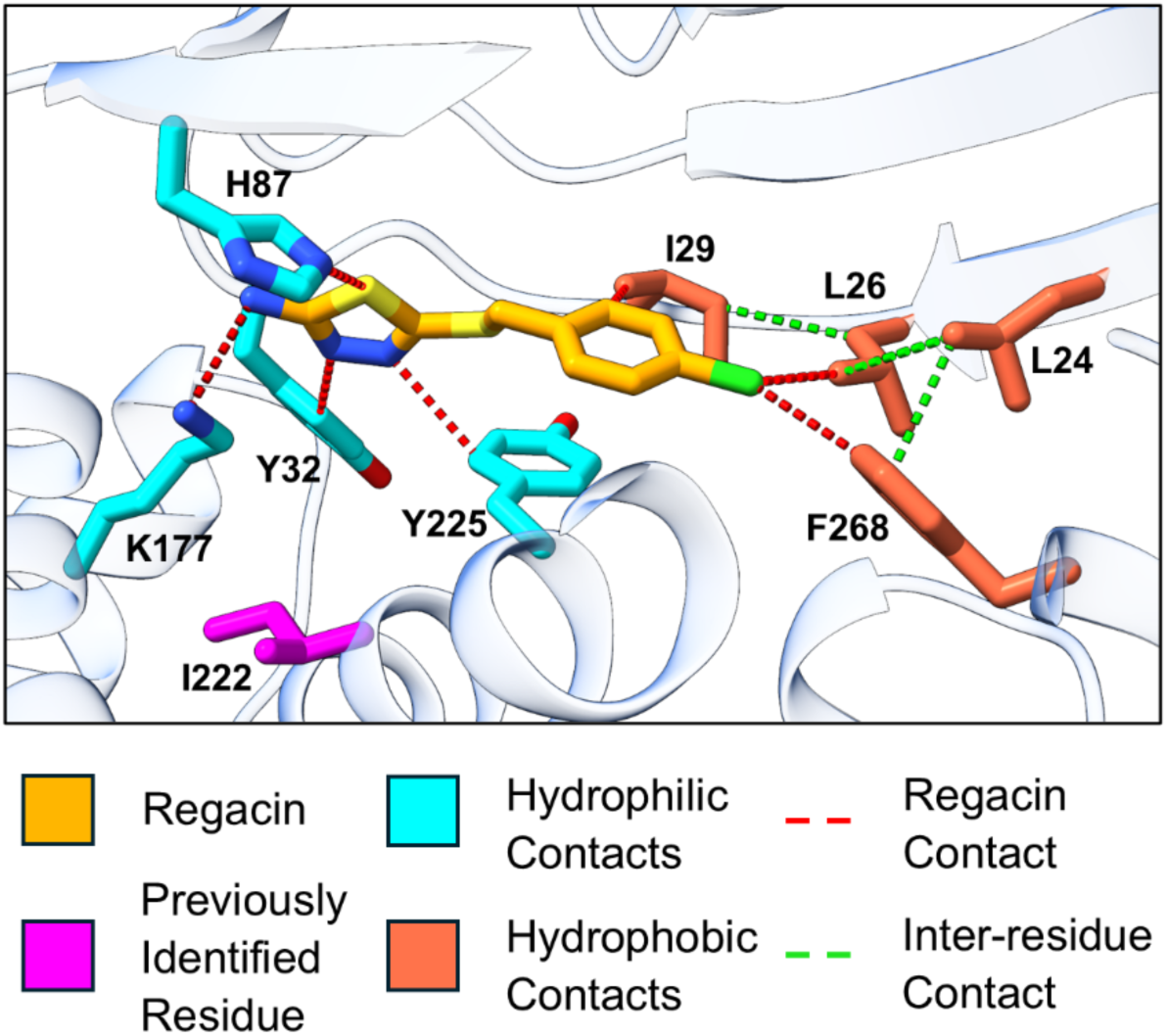
Multiple ligand binding pocket residues are important for inhibition. Residues which yield an insensitive phenotype upon mutagenesis either in this study or a previous study are indicated for regacin-RegA chain C (PDB_00001874). Polar front of pocket residues essential for inhibition are shown in cyan, hydrophobic back of pocket residues essential for inhibition are shown in orange red, and I222 identified in previous work to be implicated in inhibition is shown in magenta (Yang et al., 2013). Regacin contacts ≤4 Å are shown in red, while secondary contacts are shown in lime.

Several other notable studies have reported antivirulence molecules that inhibit virulence regulators within the AraC/XylS superfamily. Hodson et al. used a high-throughput screen to find inhibitors of Rns (CfaD). Unfortunately, they declined to identify the two inhibitors that they discovered, and the study lacked mechanistic and genetic analyses (Hodson et al., 2017). Hung et al. reported a more comprehensive high-throughput screen and analyses. Their investigations led to the discovery of virstatin (CAS 88909-96-0) which was shown to abolish ToxT-dependent expression of cholera toxin and the toxin coregulated pilus of *V. cholerae* (Hung et al., 2005). As expected, orogastric administration of virstatin protected infant mice from *V. cholerae* challenge. Like regacin, virstatin interferes with DNA binding. Unlike regacin, it appears to do so by abolishing dimerization of ToxT (Shakhnovich et al., 2007; Yang et al., 2013). As an alternative to high-throughput screens, Woodbrey et al. capitalized on the structure of ToxT with cis-palmitoleate in the binding pocket to rationally design ToxT inhibitors (Lowden et al., 2010; Woodbrey et al., 2018). They developed several inhibitory mimetics and solved the cocrystal structures of two of them showing that they bind within the ligand binding pocket. Competition binding studies strongly suggest that virstatin also binds within the pocket. Collectively, these studies demonstrate the potential therapeutic value of antivirulence drugs. High-resolution structures and in-depth binding pocket analysis of RegA, ToxT, and Rns will allow for rational drug design and accelerate AI driven drug discovery.

## Conclusion

This study defines the structural basis for inhibition of the bacterial virulence regulator RegA by the small molecule regacin. High-resolution structures of apo-and regacin-bound RegA reveal that regacin binds within a conserved ligand binding pocket in the amino-terminal domain, rather than the DNA-binding domain previously proposed as the probable site of action. Structure-guided mutagenesis further demonstrates that both polar residues near the pocket entrance and hydrophobic residues deeper within the pocket contribute to regacin sensitivity, supporting a model in which multiple interactions stabilize ligand binding and impair RegA-dependent virulence gene expression. These findings clarify how regacin inhibits RegA and establish this conserved regulatory pocket as a tractable target for the design of antivirulence therapeutics against related bacterial pathogens.

## Materials and Methods

### Construction of plasmids

pGPMRegA-myc was constructed with *regA* amplified from *C. rodentium* strain DBS100. The *regA* gene was amplified with primers 1934/1935 and NEB HiFi assembled into a pTags2 PCR fragment created with primers 1414/1419.

pAJ007 was assembled through In-Fusion cloning. The sequence encoding RegA was codon-optimized to remove rare codons, modified to include flanking sequences, purchased as a gBlock from Integrated DNA Technologies (IDT), and amplified by PCR using primers CRM27 and CRM24. The pAJ007 expression plasmid was assembled from the amplified RegA insert and linearized pCDB24 (a gift from Dr. Christopher Bahl) vector using the Takara Bio In-Fusion Cloning Kit. This assembled construct was transformed into DH5α high efficiency cells and verified by sequencing prior to transformation into BL21(DE3) cells for protein expression. The resulting pAJ007 plasmid encodes a 10xHis-SUMO-RegA fusion under T7 promoter control and confers ampicillin resistance.

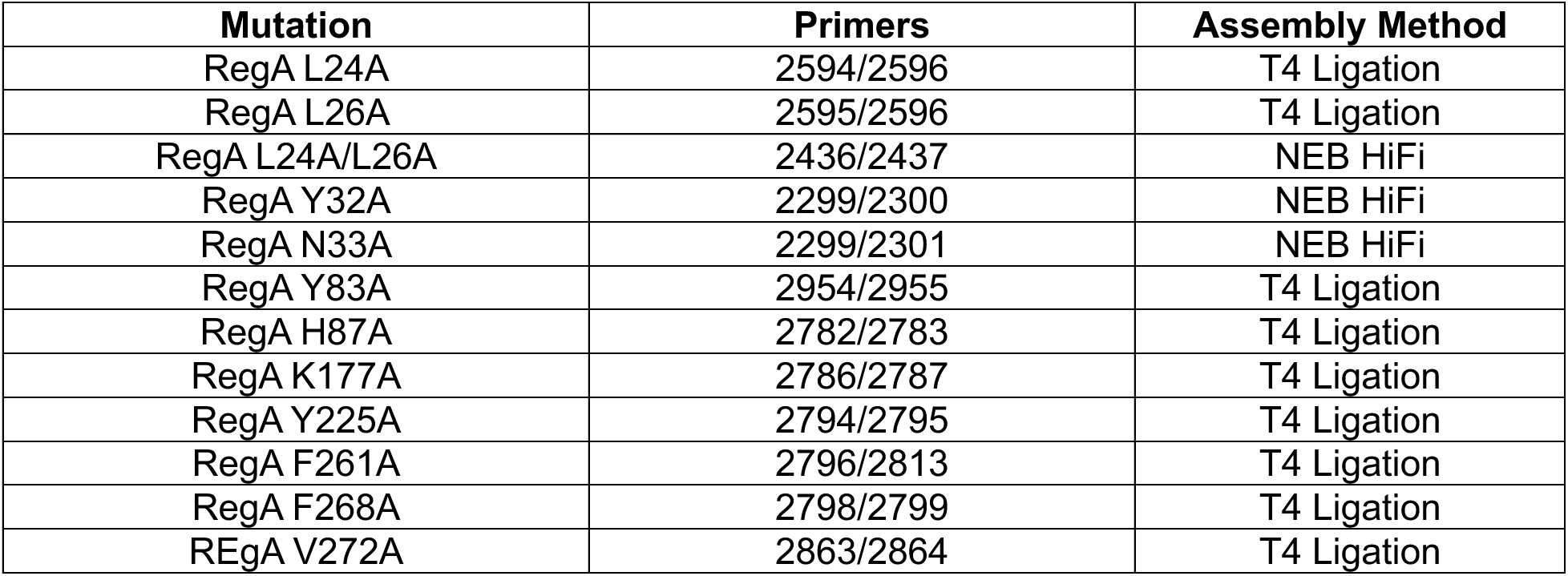

| Mutation | Primers | Assembly Method |
| --- | --- | --- |
| RegA L24A | 2594/2596 | T4 Ligation |
| RegA L26A | 2595/2596 | T4 Ligation |
| RegA L24A/L26A | 2436/2437 | NEB HiFi |
| RegA Y32A | 2299/2300 | NEB HiFi |
| RegA N33A | 2299/2301 | NEB HiFi |
| RegA Y83A | 2954/2955 | T4 Ligation |
| RegA H87A | 2782/2783 | T4 Ligation |
| RegA K177A | 2786/2787 | T4 Ligation |
| RegA Y225A | 2794/2795 | T4 Ligation |
| RegA F261A | 2796/2813 | T4 Ligation |
| RegA F268A | 2798/2799 | T4 Ligation |
| REgA V272A | 2863/2864 | T4 Ligation |

All assembled plasmids were transformed into *E. coli* DH5α and sequenced to confirm mutagenesis before being transformed into transcriptional reporter GPM1072 for analysis.

### Strain construction

Strain GPM3201 was constructed through λ RED recombination at previously described (Datta et al., 2006). A hygromycin resistance cassette was amplified from pTags-Hyg1 using primers 2450/2451. Electroporation of the PCR cassette into GPM1830/pSIM6 disrupted the *regA* gene. The resulting strain was confirmed for disruption of *regA* with primers 2456/2457, heat-cured to remove pSIM6, and labelled strain GPM3201.

Strain GPM3202 was constructed through λ RED recombination at previously described (Datta et al., 2006). A FLAG and kanamycin resistance cassette was amplified from pSUB11 using primers 1178/1179. Electroporation of the PCR cassette into GPM3201/pSIM6 added a C-terminal FLAG-tag to CexE. The resulting strain was confirmed for insertion with primers 1180/1186, heat-cured to remove pSIM6, and labelled strain GPM3202.

### Preparation of whole cell lysates for Western blot

Whole cell lysates were prepared from cultures grown aerobically at 37 °C overnight in LB broth. 500 µL of culture was pelleted at 16,000 RCF for 1 minute and resuspended in 1X SDS Loading Buffer with 400 mM β-ME before being run in SDS-PAGE gels. SDS-PAGE gels were transferred to PVDF membranes and stained with monoclonal anti-FLAG and anti-DnaK primary murine antibodies. Blots were further stained with HRP-conjugated secondary antibodies and visualized for chemiluminescence on the Invitrogen iBright 1500.

### β-galactosidase assays

RegA transcriptional activation of a RegA controlled promoter was measured via β-galactosidase assay on a Lac reporter. Strain GPM1072 or GPM1080 was transformed separately with pGPMRegA-myc or vector pTags2. To ensure proper GPMRegA-myc expression, each growth experiment started from a single colony of fresh transformation, and not frozen cells. All strains were grown aerobically at 37 °C to stationary phase in LB with 150 µg/mL ampicillin and 17.5 µg/mL of both spectinomycin and streptomycin. After 2-3 hours of growth, regacin (dissolved in DMSO) was added to a final concentration of 0.4% DMSO v/v and cultures were returned to growth conditions. All β-galactosidase experiments were assayed as previously described (Miller, 1972).

### RegA expression and purification for crystallization

RegA was expressed and purified as previously described with the following changes (Tolbert et al., 2025). Briefly, lysis was performed by sonicating on ice at 50% power for 30 seconds on then 30 seconds off for a total of 7 minutes. RegA was first captured using His-tag affinity chromatography with a Ni-NTA column, the SUMO tag was cleaved during dialysis, and removed with reverse His-tag affinity chromatography, all as previously described. The final purification was performed using a HiLoad Superdex 75 16/60 column (Cytiva) with a running buffer of 20 mM Tris pH8, 500 mM NaCl, 500 µM DTT, and 500 µM EDTA. The column was first equilibrated with 1 column volume, then RegA was loaded onto the column, and fractionated over 1 column volume. Fractions containing RegA combined and then concentrated to ∼5-6mg/mL for crystallization purposes.

### RegA crystallization

Initial crystal screening utilized matrix screens from Qiagen and Hampton in both 96-well evolution and MRC 2 drop plates (Calibre Scientific). Various ratios and drop sizes were assessed, and a 1:1 protein-to-reservoir ratio in either 2 µL or 400 nL drops yielded the most consistent crystal hits. The drops were set either by hand or using a NT8 robot and imaged with RockImager (Formulatrix) housed in the bioMT core. Initial RegA crystals were obtained in: 0.1 M Bis-Tris pH 5.7, 0.16 M NaCl and 25% PEG 3350. These initial crystals were made into a seed stock solution as described by Hampton Research. For subsequent crystallizations, the seed stock was diluted to a 1:10 ratio in the original crystallization condition, then added to ∼6 mg/mL RegA at a 2:25 seed to protein ratio. This seeded protein solution was then used for the following crystallizations. The apo-RegA structure was achieved by adding 100 mM decanoic acid in methanol to RegA for a final concentration of 0.1 mM. The best crystals were obtained by adding 1:1 ratio of the protein mixture to the crystallization solution, 200 mM KCl and 25% w/v PEG 3350, in a 400 nL drop. The reg-RegA structure was achieved by adding 100 mM regacin in DMSO to RegA reach a final concentration of 0.5 mM, then mixing 1:1 with the crystallization condition 0.1 M Bis-Tris pH 5.7, 0.16 M NaCl and 25% PEG 3350 in a 2 µL drop.

### RegA data collection, structural determination, and analysis

Diffraction data of RegA were collected at NSLS2 using either the AMX or FMX beamlines (Schneider et al., 2021, 2022). Initial data reduction was performed in autoPROC (P. Evans, 2006; P. Evans & Murshudov, 2013; Kabsch, 2010; Tickle et al., 2016; Vonrhein et al., 2011; Winn et al., 2011). For the apo-RegA and reg-RegA data sets, the isotropically scaled reflections were used. Molecular replacement was performed with PHASER by splitting the amino-terminal domain and DNA binding domain of AlphaFold RegA model and searching for them separately. Once a solution was obtained, the domains were merged into one chain using PyMOL. PHASER was used for molecular replacement (McCoy et al., 2007) with the AlphaFold 3 RegA structure prediction as the search model (Abramson et al., 2024). To phase the data, the AlphaFold 3 RegA model had to be split into the DBD and amino-terminal domains and searched for separately, then merged back together in PyMOL (Schrödinger). Automated refinement was conducted with PHENIX, with manual model building performed using COOT (Emsley et al., 2010). For the apo-RegA crystals, no density corresponding to the decanoic acid ligand was observed. For the RegA regacin crystal clear density corresponding to the ligand was observed in the binding pocket. Since there are no crystal deposition records for regacin, a regacin crystallographic information file (CIF) was built and geometry optimized in eLBOW, then placed into the structure using COOT (Moriarty et al., 2009). Both final models were deposited into the PDB database (Berman et al., 2000). Crystal data and model statistics are reported in Supplemental Table S1. Structural analysis and visualization was performed using ChimeraX (Pettersen et al., 2021).

### RegA structure visualization and analysis

ChimeraX was used to calculate RMSD values of the pairwise comparisons of the chains in the various structures using the matchmaker function with the cutoff distance set to none. A hydrophobic surface representation was created as described by the ChimeraX users’ documentation website. The Protein-Ligand Interaction Profiler (PLIP) web server was used to evaluate protein-ligand interaction types and distances outside of van der Waals contacts (Schake et al., 2025).

### Strains Table

| Name | Description | Marker | Reference |
| --- | --- | --- | --- |
| MC4100 | *E. coli* K-12 F- *araD139* $\Delta$(*argF-lac*)*U169 rpsL150* (StrR) *relA1 flhD5301 deoC1 ptsF25 rbsR* | N/A | (Casadaban, 1976) |
| GPM1072 | MC4100 *attBHK022::pCS3Lac1* | *aadA* | (Midgett et al., 2021) |
| GPM1830 | *C. rodentium* DBS100 *cexE-FLAG::kan* | *kan* | (Rivas et al., 2020) |
| GPM3201 | *C. rodentium* DBS100 *regA::hyg* | *hyg* | This study |
| GPM3202 | *C. rodentium* DBS100 *regA::hyg* *cexE-FLAG::kan* | *hyg, kan* | This study |
| BL21(DE3) | *E. coli fhuA2 [lon] ompT gal (λ DE3) [dcm] ΔhsdS* | N/A | (Studier & Moffatt, 1986) |

### Plasmid Table

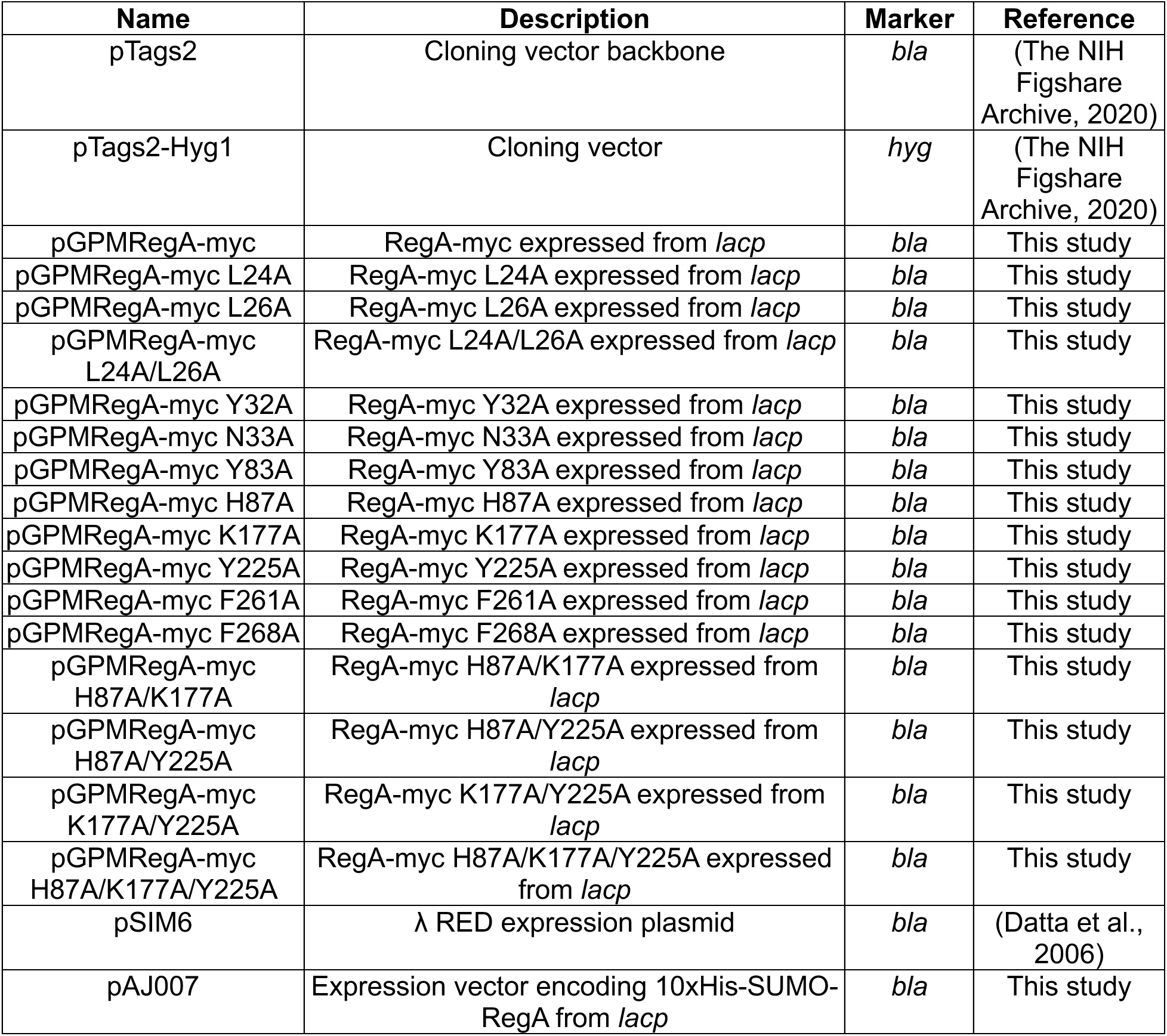

| Name | Description | Marker | Reference |
| --- | --- | --- | --- |
| pTags2 | Cloning vector backbone | *bla* | (The NIH Figshare Archive, 2020) |
| pTags2-Hyg1 | Cloning vector | *hyg* | (The NIH Figshare Archive, 2020) |
| pGPMRegA-myc | RegA-myc expressed from *lacp* | *bla* | This study |
| pGPMRegA-myc L24A | RegA-myc L24A expressed from *lacp* | *bla* | This study |
| pGPMRegA-myc L26A | RegA-myc L26A expressed from *lacp* | *bla* | This study |
| pGPMRegA-myc L24A/L26A | RegA-myc L24A/L26A expressed from *lacp* | *bla* | This study |
| pGPMRegA-myc Y32A | RegA-myc Y32A expressed from *lacp* | *bla* | This study |
| pGPMRegA-myc N33A | RegA-myc N33A expressed from *lacp* | *bla* | This study |
| pGPMRegA-myc Y83A | RegA-myc Y83A expressed from *lacp* | *bla* | This study |
| pGPMRegA-myc H87A | RegA-myc H87A expressed from *lacp* | *bla* | This study |
| pGPMRegA-myc K177A | RegA-myc K177A expressed from *lacp* | *bla* | This study |
| pGPMRegA-myc Y225A | RegA-myc Y225A expressed from *lacp* | *bla* | This study |
| pGPMRegA-myc F261A | RegA-myc F261A expressed from *lacp* | *bla* | This study |
| pGPMRegA-myc F268A | RegA-myc F268A expressed from *lacp* | *bla* | This study |
| pGPMRegA-myc H87A/K177A | RegA-myc H87A/K177A expressed from *lacp* | *bla* | This study |
| pGPMRegA-myc H87A/Y225A | RegA-myc H87A/Y225A expressed from *lacp* | *bla* | This study |
| pGPMRegA-myc K177A/Y225A | RegA-myc K177A/Y225A expressed from *lacp* | *bla* | This study |
| pGPMRegA-myc H87A/K177A/Y225A | RegA-myc H87A/K177A/Y225A expressed from *lacp* | *bla* | This study |
| pSIM6 | λ RED expression plasmid | *bla* | (Datta et al., 2006) |
| pAJ007 | Expression vector encoding 10xHis-SUMO-RegA from *lacp* | *bla* | This study |

### Oligonucleotide Table

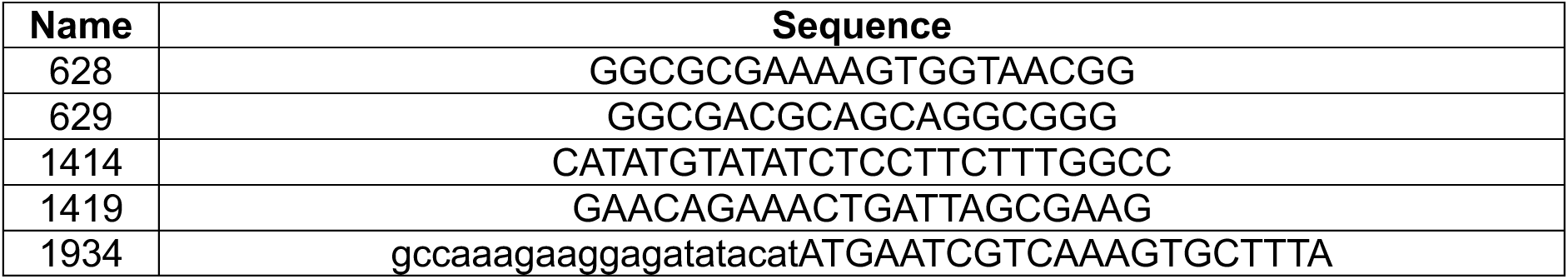
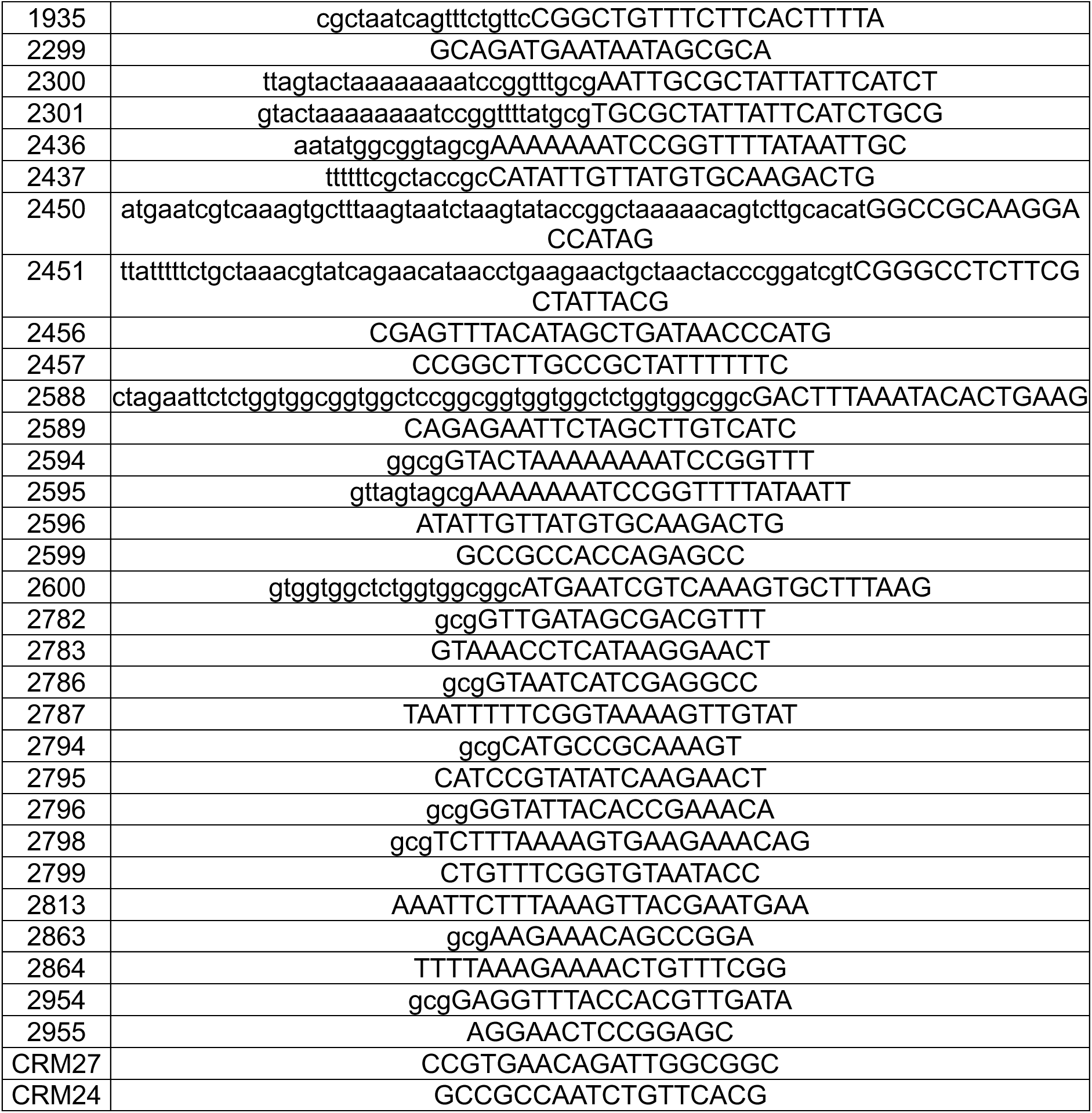

Lowercase denotes target sequences for homology-directed assembly and/or mutagenic primer sites. Italics represent core CS3 binding site.

## Acknowledgments

We would like to thank the bioMT core facility and the AMX (17-ID-1) and FMX (17-ID-2) beamline staff for making this work possible.

Research reported in this publication was supported by the National Institute of Allergy and Infectious Diseases under award number R01AI168157 (RFK and GPM) and and the National Institute of General Medical Sciences under award number P30GM165328 (bioMT) of the National Institutes of Health. The content is solely the responsibility of the authors and does not necessarily represent the official views of the National Institutes of Health.

Molecular graphics and analyses performed with UCSF ChimeraX, developed by the Resource for Biocomputing, Visualization, and Informatics at the University of California San Francisco, with support from the National Institute of General Medical Sciences under award number R01GM129325 and National Institute of Allergy and Infectious Diseases Office of Cyber Infrastructure and Computational Biology of the National Institutes of Health.

This work also used resources AMX (17-ID-1) and FMX (17-ID-2) of the National Synchrotron Light Source II, a U.S. Department of Energy Office of Science User Facility operated for the Department by Brookhaven National Laboratory under contract DESC0012704.

## Data Availability

The structures were deposited into the RCSB.org protein databank ().

## Supplemental Figures

**Table S1:** Crystallographic Statistics for RegA and Regacin-RegA Structures.

|  | **RegA** | **Regacin-RegA** |
| --- | --- | --- |
| **Wavelength** | 0.92010 | 0.92010 |
| **Resolution range** | 33.34 - 2.69 (2.74 - 2.69) | 48.15 - 2.55 (2.63 - 2.55) |
| **Space group** | P 1 21 1 | P 1 21 1 |
| **Unit cell a b c (Å)** | 68.293 98.569 86.170 | 68.154 98.842 86.255 |
| **Unit cell $\alpha$ $\beta$ $\lambda$ (°)** | 90.00 100.23 90.00 | 90.00 101.13 90.00 |
| **Total reflections** | 109358 (10033) | 133789 (13848) |
| **Unique reflections** | 31082 (2826) | 42426 (4419) |
| **Multiplicity** | 3.5 (3.6) | 3.2 (3.1) |
| **Completeness (%)** | 99.58 (99.08) | 94.08 (99.45) |
| **Mean I/sigma(I)** | 5.44 (0.81) | 4.34 (0.82) |
| **R-merge** | 0.1285 (1.248) | 0.1092 (1.315) |
| **R-meas** | 0.1522 (1.473) | 0.1317 (1.585) |
| **R-pim** | 0.08074 (0.7779) | 0.07264 (0.8742) |
| **CC1/2** | 0.994 (0.439) | 0.991 (0.346) |
| **Reflections used in refinement** | 31043 (2805) | 34513 (3613) |
| **Reflections used for R-free** | 1543 (118) | 1689 (189) |
| **R-work** | 0.2165 (0.3457) | 0.2141 (0.3106) |
| **R-free** | 0.2514 (0.3662) | 0.2630 (0.3510) |
| **Number of non-hydrogen atoms** | 8143 | 8231 |
| **macromolecules** | 8096 | 8120 |
| **ligands** | 1 | 35 |
| **solvent** | 46 | 76 |
| **Protein residues** | 1001 | 1004 |
| **RMS (bonds)** | 0.003 | 0.003 |
| **RMS (angles)** | 0.62 | 0.61 |
| **Ramachandran favored (%)** | 97.77 | 96.77 |
| **Ramachandran allowed (%)** | 2.23 | 3.12 |
| **Ramachandran outliers (%)** | 0 | 0.1 |
| **Average B-factor** | 75.05 | 60.53 |
| **macromolecules** | 75.12 | 60.51 |
| **ligands** | 68.4 | 77.71 |
| **solvent** | 62.2 | 54.92 |

**Figure S1:**
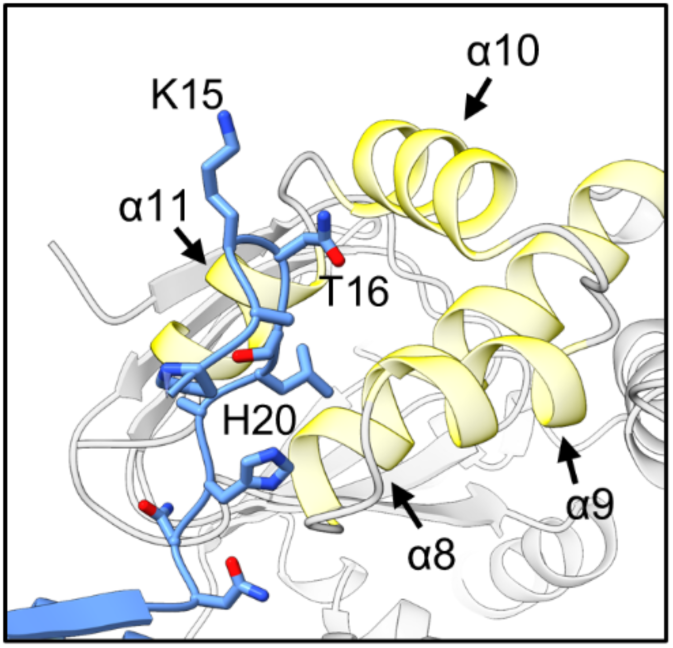
Regacin-RegA chain A amino-terminus (cornflower blue) interacting with the symmetry mate DBD helices α8, α9, α10, and α11 (yellow).

**Figure S2:**
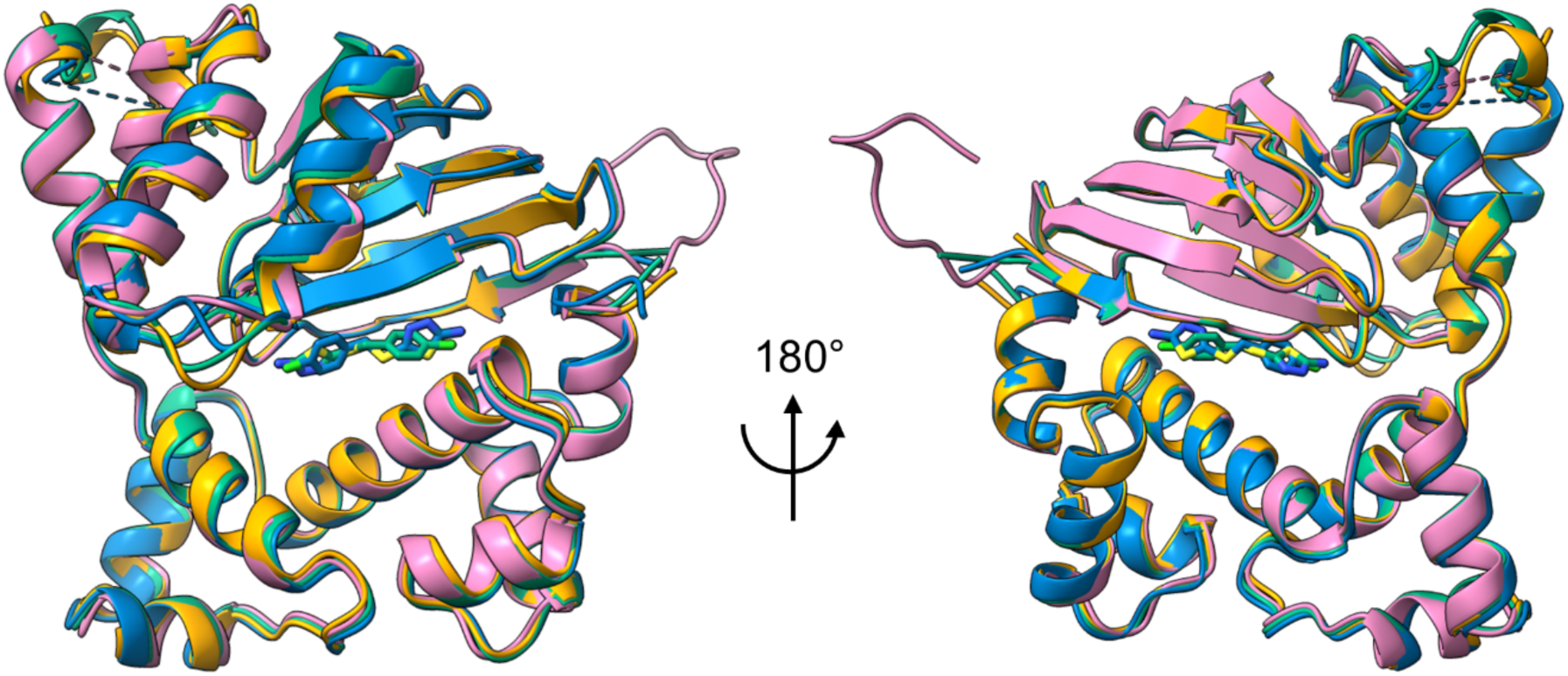
Each chain in the regacin-RegA asymmetric unit is aligned, and they are nearly superimposable. Chain A is shown in pink, B is orange, C is green and D is blue. The loop regions display variable conformations and are unresolved in some chains (chain breaks shown as dashed lines.) The regacins from chain C and D align very well within the pocket but are in different orientations.

**Figure S3:**
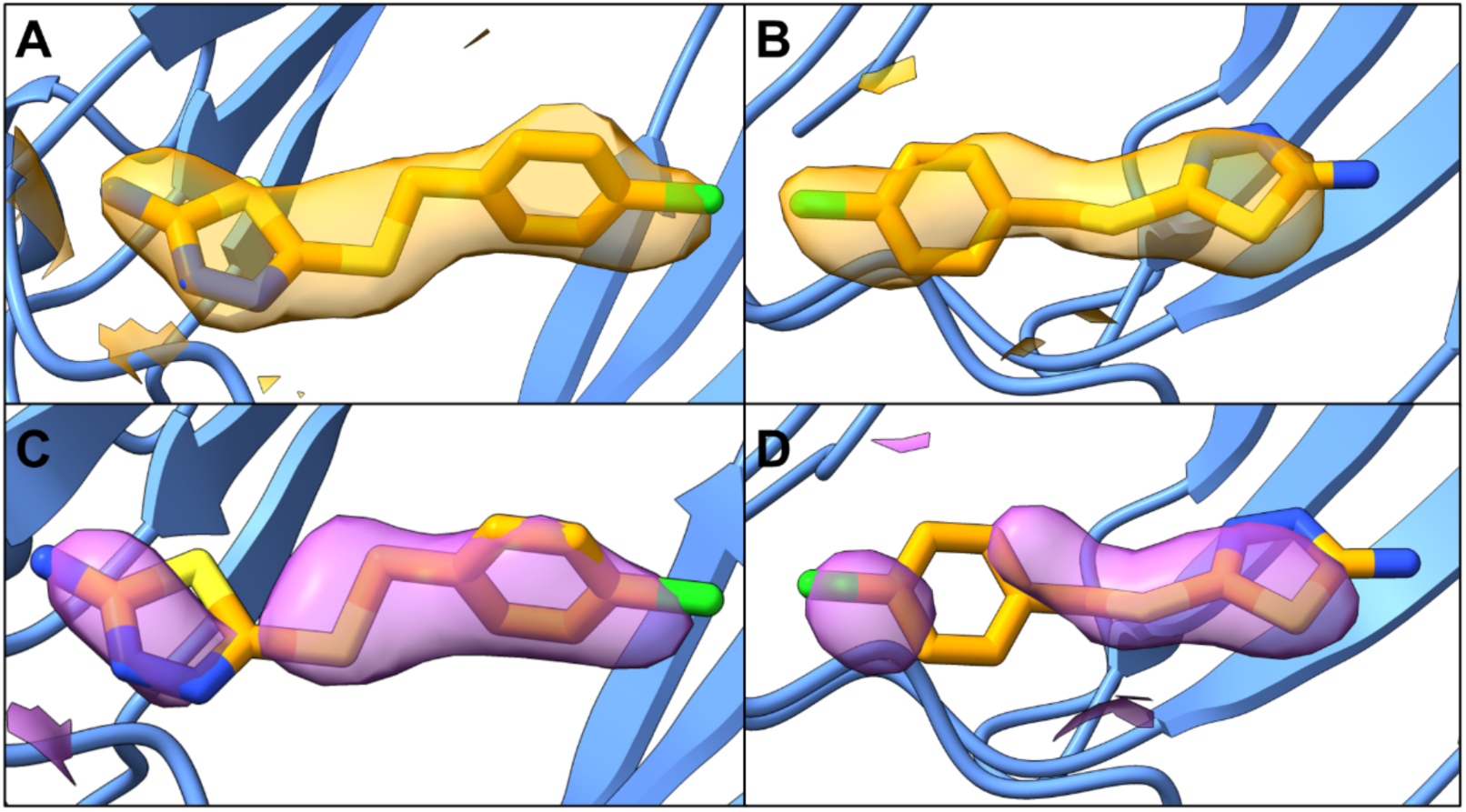
**A)** Regacin-RegA 2Fo-Fc electron density map (orange) contoured at 1 σ (RMSD) and displayed within a 2.5 Å radius of the regacin ligand bound in the pocket of chain B. **B)** Same map as in panel A, but showing the density around the regacin ligand in the pocket of chain D. **C)** Regacin-RegA composite-omi 2mFo-DFc map (purple) contoured at 1 σ (RMSD) and displayed within a 2.5 Å radius of the regacin ligand in the pocket of chain B. **D)** Same composite-omit map as in panel C, but for the regacin ligand bound in the pocket of chain D.

## Notes

### Competing Interest Statement

The authors have declared no competing interest.

### Summary of Updates

Needed to an author to the author list.

